# Gut Microbiota Modulates and Predicts Disease Severity in Experimental Pemphigoid Disease

**DOI:** 10.1101/2025.10.01.679837

**Authors:** Xiaolin Liu, Sabrina Patzelt, Yue Ma, Aleksa Cepic, Hasan-Onur Dikmen, Katja Bieber, Mathieu Groussin, Mathilde Poyet, Saleh Ibrahim, Enno Schmidt, John F. Baines

## Abstract

Pemphigoid diseases (PD) are autoimmune blistering diseases with reported alterations in skin and gut microbiota, though their causal contribution to disease pathophysiology remains unclear. Using a passive model of bullous pemphigoid-like epidermolysis bullosa acquisita (BP-like EBA), we compared C57BL/6J mice from two sources (in-house vs. Charles River) that differed in their baseline microbiota. Charles River mice developed significantly less severe disease. Co-housing led to partial homogenization of the gut microbiota, driven by asymmetric transfer of taxa from Charles River to in-house mice, which corresponded with reduced disease severity in the latter. The skin microbiota, however, showed limited exchange. Disease severity was inversely associated with gut microbial alpha diversity, with protective taxa such as *Lactobacillus intestinalis* and *Parabacteroides distasonis* enriched in Charles River mice, while pro-inflammatory taxa including *Turicimonas muris* and *Muribaculum intestinale* were enriched in in-house mice. A machine learning model further identified a gut taxon most closely matching the candidate genus *Scatocola* as a strong negative predictor of disease severity. Through experimental exposure and uptake of variable gut microbiota, these findings suggest a direct role of gut microbiota in mediating the severity of the experimental BP-like EBA and highlight the potential of microbiota-based strategies for therapeutic intervention in PD.

## Introduction

Pemphigoid diseases (PD) are a group of autoimmune blistering disorders (AIBD) that primarily affect the skin and surface-close mucous membranes, characterized by the formation of blisters and erosions resulting from autoantibodies targeting structural proteins of the muco-cutaneous basal membrane zone (BMZ) (1). The most frequent PD is bullous pemphigoid (BP), which typically affects the elderly (2, 3). Clinically, BP presents with tense blisters, erosions, erythematous macules and plaques and severe pruritus. BP is associated with increased morbidity, decreased quality of life, and increased mortality (3, 4). Autoantibodies target BP180 (also termed collagen type XVII; Col17) in nearly all patients, and BP230 in about half of them (1). Binding of autoantibodies to Col17 and BP230 results in a mainly type 2 inflammatory response including the activation of complement along the muco-cutaneous BMZ, the influx of inflammatory cells such as eosinophils, T lymphocytes, macrophages and neutrophils in the upper dermis. The release of reactive oxygen species and specific proteases from these cells finally leads to subepidermal splitting (3). While certain drugs have been described as inducers of BP (5), in most patients the trigger factors that lead to the break of tolerance against Col17 and BP230 in this elderly patient population are unknown. Thus, the need to further understand the pathogenesis and factors associated with the initiation and progression of PD is instrumental to improve the lives of PD patients. Recently, in a large-scale prospective study with 228 BP patients, we described alterations in the skin microbiota, including the enrichment of *Staphylococcus aureus* as an inflammation-promoting species in BP patients (6). Further, at the time of diagnosis, the skin microbiota was also found to be significantly altered at non-lesional skin sites in BP patients compared to site-, age-, and sex-matched controls (6). These differences suggest skin microbiota as a potential trigger factor in BP. Furthermore, BP lesions were shown to contain significantly more toxic shock syndrome toxin-1 superantigen-producing *Staphylococcus aureus* compared to both non-lesional BP skin and skin of control patients (7).

In addition to the potential role of skin microbiota, increasing evidence highlights the significance of a gut-skin axis, linking gut microbiota and skin health, in inflammatory dermatoses such as atopic dermatitis, psoriasis, chronic spontaneous urticaria and, more recently, in pemphigus and pemphigoid diseases (8–24). Gut dysbiosis can influence skin health by e.g. triggering immunological responses through microbial metabolites such as SCFA and GABA (6, 25). In our recent study of the gut microbiome in 66 BP patients and their age- and sex-matched controls, decreased alpha-diversity and an overall altered gut microbial community was observed, together with reduced *Faecalibacterium prausnitzii* and a greater abundance of *Flavonifractor plautii* (22). Thus, importantly, the gut microbiome in BP, an inflammatory skin disorder, displays changes in the gut microbiome that resemble the hallmark dysbiotic signatures found in diseases such as inflammatory bowel disease (22, 26).

Several mouse models have been developed for PD, including BP and BP-like epidermolysis bullosa acquisita (EBA), a PD characterized by autoimmunity against Col7. These PD mouse models reflect major clinical and immunopathological features of the human diseases and are induced by either the repeated injections of rabbit IgG against Col 17/ Col7 (passive models), or immunization of mice with recombinant fragments of murine Col 17/ Col7 (active models) (27–30). Using these PD mouse models, we previously showed that mice that did not develop clinical disease, although serum anti-Col7 IgG and tissue-bound IgG and C3 were present, revealed a significantly higher richness and distinctly clustered diversity of their skin microbiota compared to diseased animals (31). In the same model, genetic loci were identified that contributed to skin microbiota variability (32).

The specific role of gut and skin microbiota in PD pathophysiology remains however poorly characterized, and experiments demonstrating microbial modification of disease severity are lacking. Here, we leveraged the fact that animal facility effects can lead to distinct microbial community differences, even between mice belonging to the same genetic background, as we previously demonstrated for C57BL/6J mice (33). In preliminary experiments, we observed C57BL/6J mice that differ according to their commercial provider and breeding origin (in-house vs. Charles River) also display significant differences in PD disease score. We thus hypothesized that underlying differences in gut and/or skin microbiota may be responsible. Through co-housing experiments, we show that homogenization of the gut microbiota and the transfer of individual pro-vs. anti-inflammatory gut taxa significantly associates with disease severity, whereas comparatively little transfer and disease association was observed for the skin microbiota. These results highlight the pathogenetic relevance of the gut-skin axis in PD and may open new therapeutic avenues by modulating gut microbial composition.

## Results

## 1. Response to PD induction and baseline gut and skin microbiota

In preliminary experiments, we observed differences in PD disease score between mice derived from in-house breeding at the University of Lübeck (UL) compared to those obtained directly obtain from Charles River (CR), whereby UL mice displayed significantly elevated disease scores (**Fig. S1**). To investigate whether (i) differences in the underlying gut and/or skin microbiota may be responsible for these observations and (ii) effects on disease severity could be transferred between mice, we performed co-housing experiments involving a total of 90 mice (**Fig. 1**). These comprised 30 UL mice, 30 CR mice, and 30 mice that underwent 28 days of co-housing prior to disease induction (Mix groups). The Mix groups were further divided according to their origin, i.e. UL mice that were housed together with CR mice (Mix_UL; n=16), and CR mice that were housed together with UL mice (Mix_CR; n=14). Within each group, the mice were randomly and equally divided into 2 further groups: one injected with anti-COL7 IgG (PD group, n = 15 for UL and CR, n = 8 for Mix_UL, n = 7 for Mix_CR) to induce BP-like EBA, and the other receiving IgG isolated from healthy rabbits as controls (normal rabbit IgG, NR groups).

**Fig. 1.**
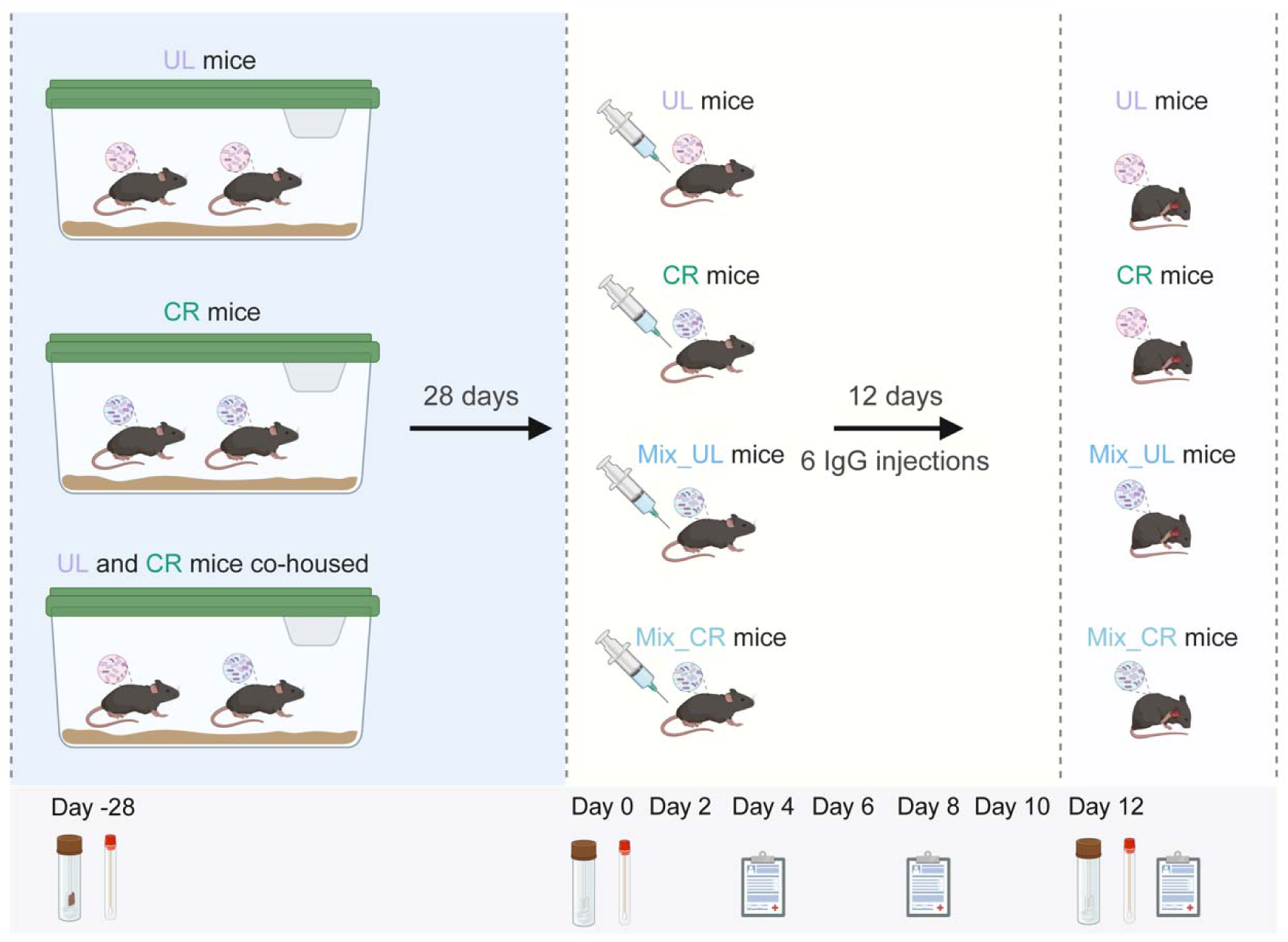
Experimental workflow of investigation on gut and skin microbiota in the PD mouse model. UL refers to mice from mouse facility of university of Lübeck (n = 30); CR indicates mice from Charles River breeding (n = 30); Mix groups contain the 30 co-housed mice, 16 of which are from UL (Mix_UL) and 14 of which are from CR (Mix_CR) source. Those mice were injected with either anti-COL7 IgG (PD group, n = 15 for each source) to induce BP-like EBA model, or normal rabbit IgG as controls (NR group, n = 15 for each source).

After 12 days, disease severity varied significantly among mouse groups from different sources (**Fig. 2A, Supplementary Table 1**). Specifically, in line with preliminary experiments, the UL mice again developed more severe skin lesions than CR mice (p = 0.00034), as well as Mix_CR mice (p = 0.0047). Intriguingly, the co-housed mice, particularly the Mix_UL group, show an intermediate level of disease severity, with Mix_UL trending higher than Mix_CR, marginally significantly lower than UL (p = 0.08), and higher than the CR mice (p = 0.06) (**Fig. 2A**). Overall, severity decreases in the following order for all mice on day 12: UL > Mix_UL > Mix_CR > CR. Moreover, the disease scores over the course of disease development (days 4 and 8) were also collected and used to assess the rates of disease progression, which are indicated by the slopes in a mixed linear model with disease severity as outcomes, and time, group and interaction of group and time as fixed effects (**Fig. 2B**). This reveals UL mice to have the fastest progression rate, significantly higher than co-housed (Mix_UL: p.adj = 0.04; Mix_CR: p.adj = 0.0048)- and CR mice (p.adj = 2.5e-06). The progression shows a similar trend as the final disease severity on day 12, and ranks as follows: UL > Mix_UL > Mix_CR > CR, with Mix_UL also significantly higher than CR (p.adj = 0.04).

**Fig. 2.**
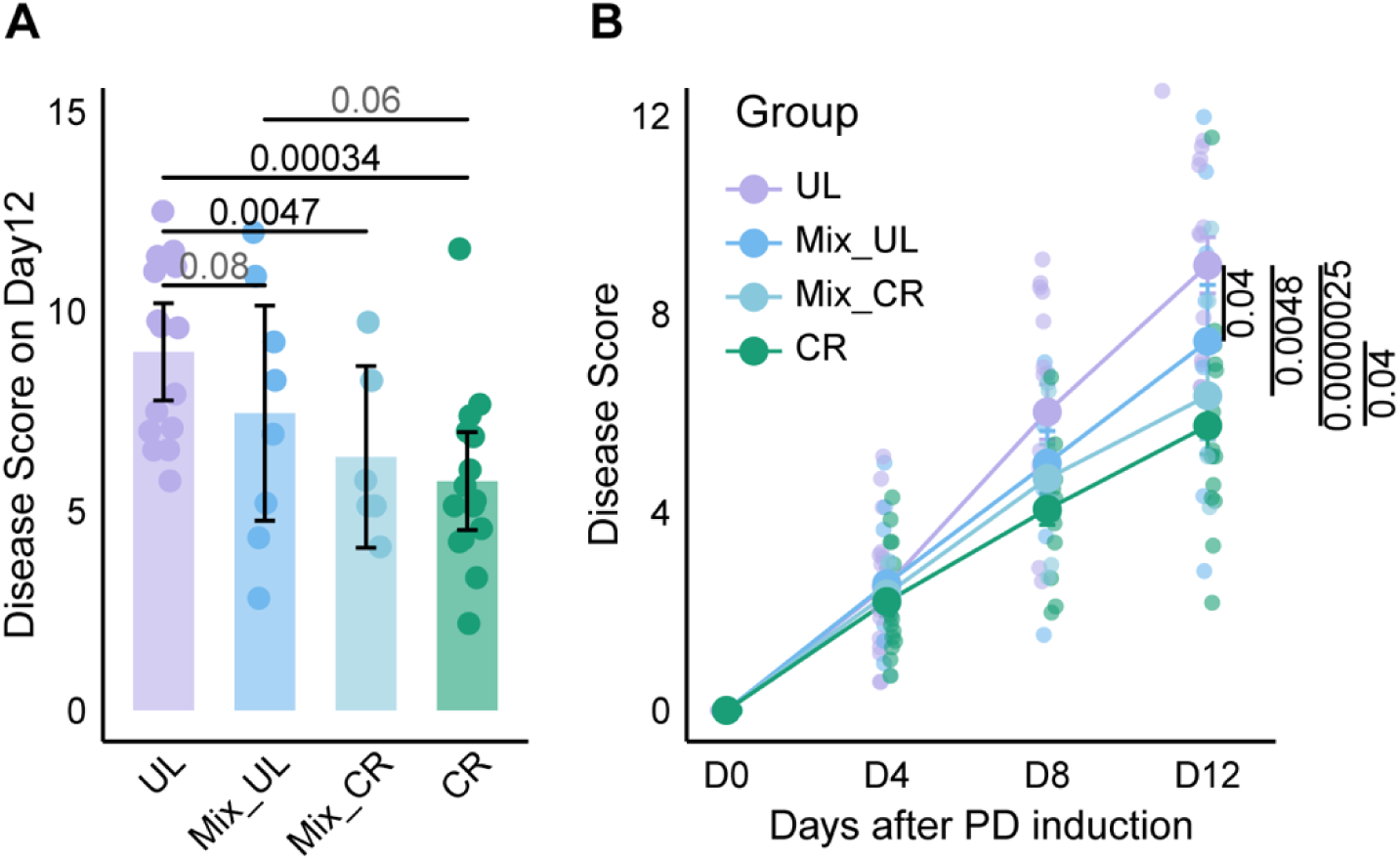
Evaluation of disease progression and severity across PD mice groups. (A) The disease score for mice from CR and UL sources and co-housed UL (Mix_UL) and CR (Mix_CR) mice on day 12. The scores were compared with t test. (B) The disease development over the 12-day course of PD induction. The disease progression across PD mice groups were assessed using a linear mixed-effects model, with disease score as the outcome, group, time, and their interaction as fixed effects, while accounting for individual mouse variability. Post-hoc comparisons of slopes were performed, with significance adjusted via the Benjamini-Hochberg (BH) procedure. The different colours represent different groups.

To evaluate the potential role of gut and skin microbiota, we performed 16S rRNA gene amplicon-based sequence analysis of gut and skin samples collected at three timepoints: day -28 (the starting day of co-housing for the 30 mice in the Mix groups), day 0 (the starting timepoint of disease induction), and day 12 (the endpoint of the experiment) (**Fig. 1**). To first focus on the microbial variation present at the baseline, we analyzed the microbiota samples from day 0. For gut microbiota, we observe significant differences in both alpha (Shannon index) and beta diversity between groups. The Shannon index is significantly higher in CR mice compared to UL mice (p = 0.029), with Mix mice displaying intermediate values (**Fig. 3A**). Beta diversity analysis also reveals significant differences among the four groups, with a gradual transition in gut microbiota composition from UL to CR mice (**Fig. 3B**). Moreover, the gut microbiota of Mix_UL and Mix_CR mice exhibits higher similarity compared to that of UL and CR mice, suggesting that co-housing for 28 days led to partial homogenization of the gut microbiota. Similarly, the skin microbiota displays significantly higher Shannon index for CR mice compared to UL (p = 0.0016) as well as Mix_UL mice (p = 0.001). The Mix_CR mice also show a significantly higher Shannon index compared to UL (p = 0.0018) and Mix_UL mice (p = 0.0045), with similar differential trends for the Chao1 index (**Fig. 3C**). Beta diversity analysis further confirmed significant differences in skin microbiota composition among the groups. However, unlike the gut microbiota, the skin microbiota of mice from the same original source (UL/Mix_UL and CR/Mix_CR) retained greater similarity, indicating that the skin microbiota was less influenced by co-housing compared to the gut microbiota (**Fig. 3D**).

**Fig. 3.**
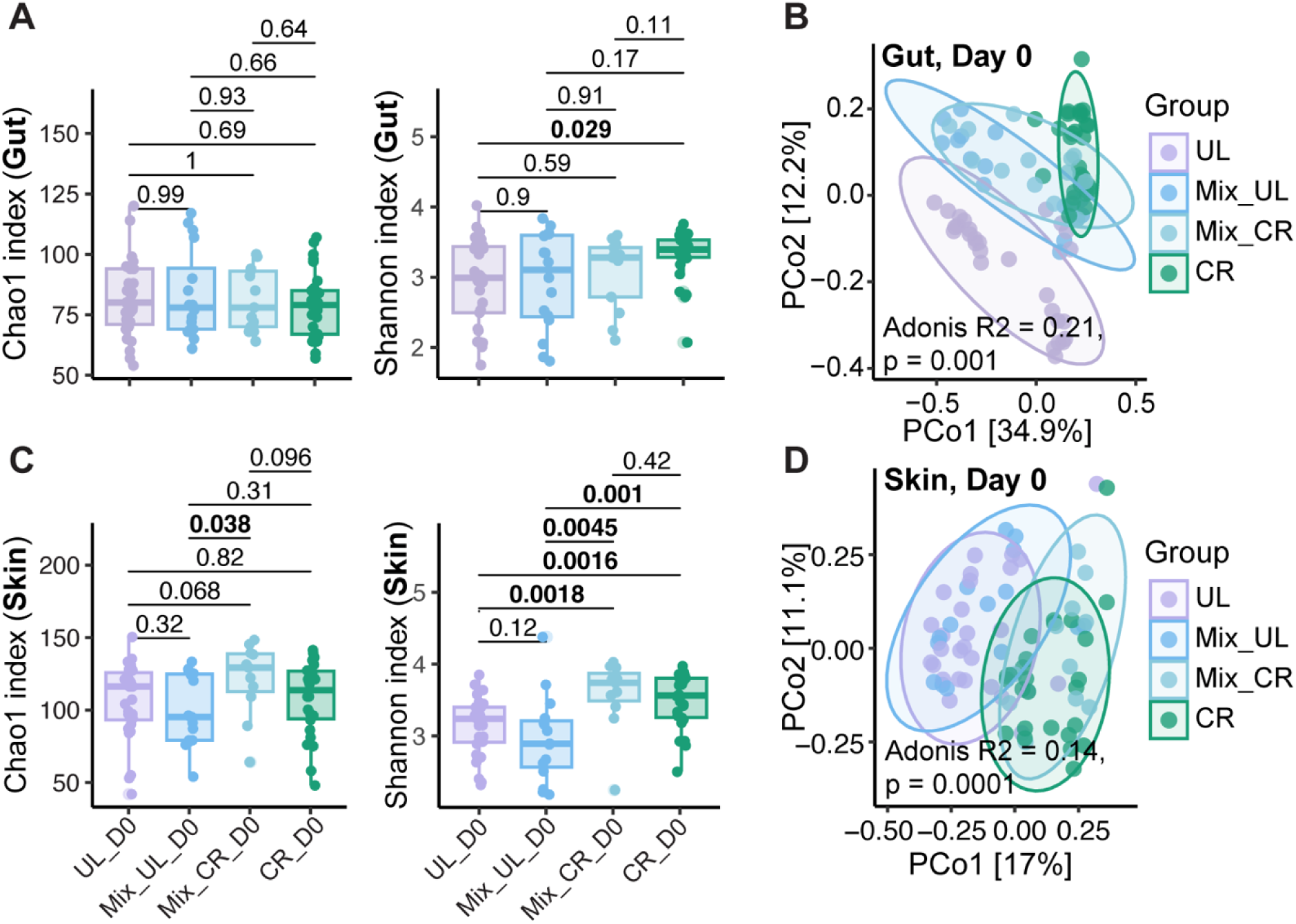
Alpha and beta diversity of gut and skin microbiota before PD induction. (A) Chao1 and Shannon indices are representing the richness and evenness of gut microbiota of UL, co-housed UL, co-housed CR and CR mice; (B) The PCoA based on Bray-Curtis dissimilarity shows the distinct clusters of gut microbiota of mice from 4 groups; (C) Chao1 and Shannon indices are representing the richness and evenness of skin microbiota mice from 4 groups; (B) The PCoA (principal coordinates analysis) based on Bray-Curtis dissimilarity shows the distinct clusters of skin microbiota of mice from 4 groups. The alpha indices were compared with Wilcoxon test. Beta diversity was analysed using PERMANOVA with 9,999 permutations.

To explore potentially protective and harmful microbes associated with disease onset and progression, we identified differentially abundant gut and skin Operational Taxonomic Units (OTUs) clustered at 99% identity prior to PD induction (day 0). Specifically, a total of 82 gut OTUs and 33 skin OTUs were found to differ among groups before disease induction (p.adj < 0.05, LDA > 2; Linear Discriminant Analysis Effect Size (LEfSe)) (**Fig. S2A-B,** only LDA >3.5 were displayed; **Supplementary Table 2**). In the gut, 50 OTUs are enriched in CR mice, compared to only 21 being enriched in UL mice, which is consistent with the higher alpha diversity observed in CR compared to UL mice. Additionally, 6 OTUs are enriched in co-housed CR mice (Mix_CR), and 4 are enriched in co-housed UL mice (Mix_UL). Notably, gOTU_2 (matching *Turicimonas muris* in NCBI, E-value = 1e-149, identity = 100%) is enriched in UL mice. This species was previously reported to be positively associated with pro-inflammatory cytokines such as TNF-alpha and IL-6 and negatively correlated with fecal butyrate levels (34). Similarly, gOTU_36, matching *Muribaculum intestinale* (NCBI, E-value = 7e-153, identity = 100%), is also enriched in UL mice, and was linked to the stimulation of pro-inflammatory cytokines, including TNF-alpha, IL-6, and IL-23 (35).

In contrast, a number of potentially protective gut microbes are enriched in CR mice. These include: (i) gOTU_15, most closely matching *Prevotellaceae_NK3B31_group*, reported to be positively associated with levels of acetic acid, propionic acid and total SCFAs in the colonic digesta, and negatively associated with the severity of dextran sulfate sodium (DSS)-induced colitis (36, 37); (ii) gOTU_53, identified as *Lactobacillus intestinalis* (NCBI, E-value = 3e-158, identity = 100%), reported to exert protective effects against DSS-induced colitis in mice (38); (iii) gOTU_93, matching *Parabacteroides distasonis* (NCBI, E-value = 2e-146, identity = 100%), reported to be reduced in rheumatoid arthritis and alleviates arthritis symptoms in mice by restoring the Th17/Treg balance (39); (iv) gOTU_24, most closely matching *Lachnospiraceae_NK4A136_group*, reported to be an SCFA-producer and negatively correlated with multiple metabolic- and inflammatory disorders (40); and (v) gOTU_215, belonging to the genus *Roseburia*, and known to play critical roles in anti-inflammatory processes, maintaining intestinal barrier integrity, and regulating intestinal hormone levels (41, 42) (**Fig. S1A**). In the skin microbiota, 16 OTUs are abundant in CR mice, with seven enriched in UL, nine in Mix_CR, and only one in Mix_UL. Notably, sOTU_1058, most closely matching *Staphylococcus epidermidis* (NCBI, E-value = 4e-150, identity = 100%), is enriched in CR mice. *S. epidermidis* is known to downregulate pro-inflammatory genes (e.g., TNFα, IL1β, IL6), enhance resistance to *S. aureus* colonization, and reduce atopic dermatitis-like skin inflammation in mice (43).

### 2. The influence of co-housing on the gut and skin microbiota

As described above, the gut microbiota displays evidence of homogenization after four weeks of co-housing (**Fig. 3B**), while the skin microbiota appears to have remained largely distinct (**Fig. 3D**). Given this higher similarity of gut microbiota composition between co-housed Mix_UL and Mix_CR at day 0, we next compared the gut microbiota before (day -28; hereafter “Dm28”)- and after co-housing (day 0; hereafter “D0”) to investigate the impact of co-housing on the microbial composition in the gut of the co-housed mice. At Dm28, significant differences in both alpha diversity (Chao1 and Shannon indices) and beta diversity are observed between the UL and CR mice destined for co-housing (i.e. Mix_CR and M_UL). The Mix_CR mice display significantly higher richness (Chao1 index, p = 0.020) and evenness (Shannon index, p = 0.0016), as well as a significantly distinct cluster in composition compared to the Mix_UL group. However, these differences in diversity metrics are no longer significant at D0. The Chao1 and Shannon indices of Mix_UL mice increased and the beta diversity clusters of Mix_CR and Mix_UL disappeared (**Fig. 4A-C**), suggesting that microbial transfer during co-housing enhanced microbial diversity in the Mix_UL group.

**Fig. 4.**
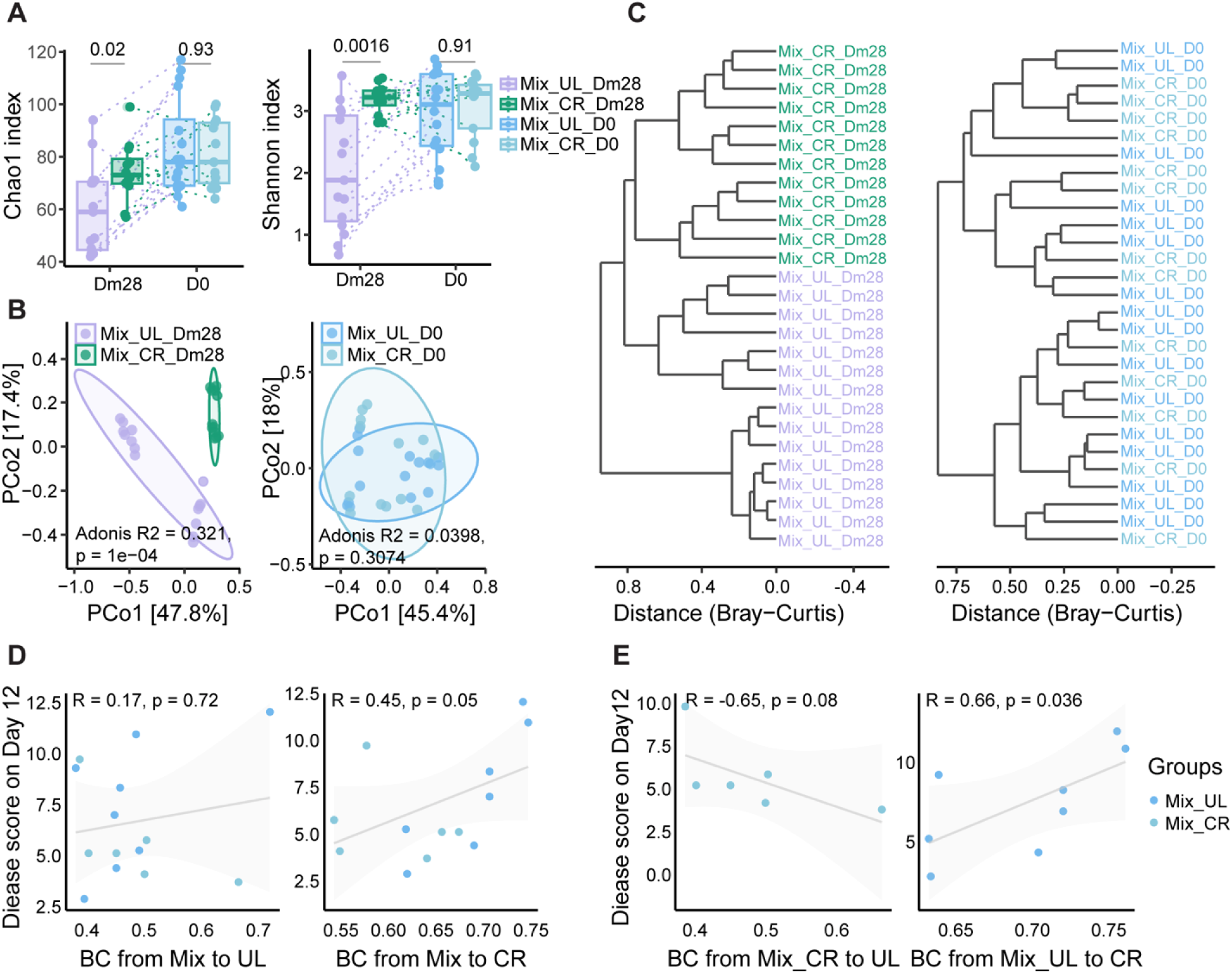
Evaluation of the potential effects of co-housing on the gut microbiota composition and disease development. (A) Chao1 and Shannon indices of gut microbiota in 30 co-housed mice before (Dm28) and after (D0) co-housing; (B) The PCoA plots of gut microbiota of co-housed UL and CR mice before (Dm28) and after (D0) co-housing; (C) The hierarchical clustering (hcluster) plots based on Bray-curtis distance are indicating the similarity of gut microbial community of individual mouse before and after co-housing. (D) The Correlation between the disease outcome on day 12 and the (BC) Bray-Curits distance from co-housed mice to UL or CR mice on day 0. (E) The correlation between the disease outcome on day 12 and BC distance between co-housed CR mice to the UL group (left) and co-housed UL mice to the CR group (right). The alpha indices were compared with Wilcoxon test. Beta diversity was analysed using PERMANOVA with 9,999 permutations.

Given the observed differences in disease severity across different groups and the potential protective role of the gut microbiota of CR mice, which is indicated by the reduced disease severity and progression rate in Mix_UL mice compared to UL mice, along with the partial homogeneity of the gut microbiota of Mix_UL and Mix_CR mice after co-housing, we next investigated the correlation between the dissimilarity (Bray-curtis distance on day 0) between co-housed mice and UL and CR groups and disease outcome (disease score on day 12). Importantly, we observe that the greater the dissimilarity between the gut microbiota of co-housed mice and CR mice before PD induction, the higher the disease severity (corr = 0.45, p = 0.05). The dissimilarity between co-housed mice and UL is however not significantly associated with disease scores (**Fig. 4D)**. Furthermore, the correlation between dissimilarity and disease severity was separately calculated for the Mix_CR and UL group, and for the Mix_UL and CR. These results indicate that the greater the dissimilarity in gut microbiota between Mix_CR mice and UL, the lower the disease score (marginally significant, corr = -0.65, p = 0.08). In contrast, the gut microbiota dissimilarity between Mix_UL mice and CR is significantly positively correlated with disease severity (corr = 0.66, p = 0.036). These results thus suggest that protective microbial features present in the CR gut microbiota were transferred to Mix_UL during co-housing.

To identify microbes potentially exchanged during co-housing, we next investigated differential gut OTUs before (Dm28) and after co-housing (D0). A total of 49 OTUs significantly differ before co-housing, with only 1 remaining significant afterward (**Fig. S3**, only OTUs with LDA > 5 are showed for Dm28; **Supplementary Table 3**). Among the pre-cohousing differences, gOTU_3, most closely matching *Lactobacillus johnsonii* (NCBI, E value = 5e-162, identity = 100%), and gOTU_7, matching genus *Alistipes*, were enriched in Mix_CR mice. *L. johnsonii* is a well-known beneficial gut microbe with anti-inflammatory properties (44–46), whereas *Alistipes* is reported to mitigate colitis in mice (47). Other potentially beneficial taxa enriched in this group include gOTU_15 (genus *Prevotellaceae_NK3B31_group*), gOTU_7 (genus *Alistipes*) and gOTU_53 (species *Lactobacillus intestinalis*) (**Fig. S3A**). In contrast, gOTU_2, enriched in UL mice prior to disease induction, is also enriched in the Mix_UL group before co-housing. Following co-housing, only gOTU_98 (genus *Ruminococcus*) remains differentially abundant, with higher levels in Mix_CR mice (**Fig. S3B**).

To complement this analysis, we next applied Sourcetracker, a Bayesian-based algorithm, to quantify the contribution of microbial communities from potential sources to recipient (“sink”) communities (48). Microbiota profiles collected after co-housing (Mix_UL_D0 and Mix_CR_D0) were designated as sinks, while pre-cohousing profiles (Mix_UL_Dm28 and Mix_CR_Dm28) served as sources, enabling assessment of microbial transfer between co-housed mice. In total, 46 gut OTUs were significantly transferred from Mix_CR to Mix_UL, compared with 16 transferred in the opposite direction (p.adj < 0.05) (**Fig. 5A-B; Supplementary Table 4**). Consistent with the differential abundance analysis, several potentially beneficial OTUs, including gOTU_3, gOTU_4, gOTU_7, gOTU_15, and gOTU_53, were transferred from Mix_CR to Mix_UL, whereas gOTU_2 was transferred from Mix_UL to Mix_CR. This asymmetric transfer pattern indicates that the CR gut microbiota exerted a stronger influence on the UL gut microbiota during co-housing.

**Fig. 5.**
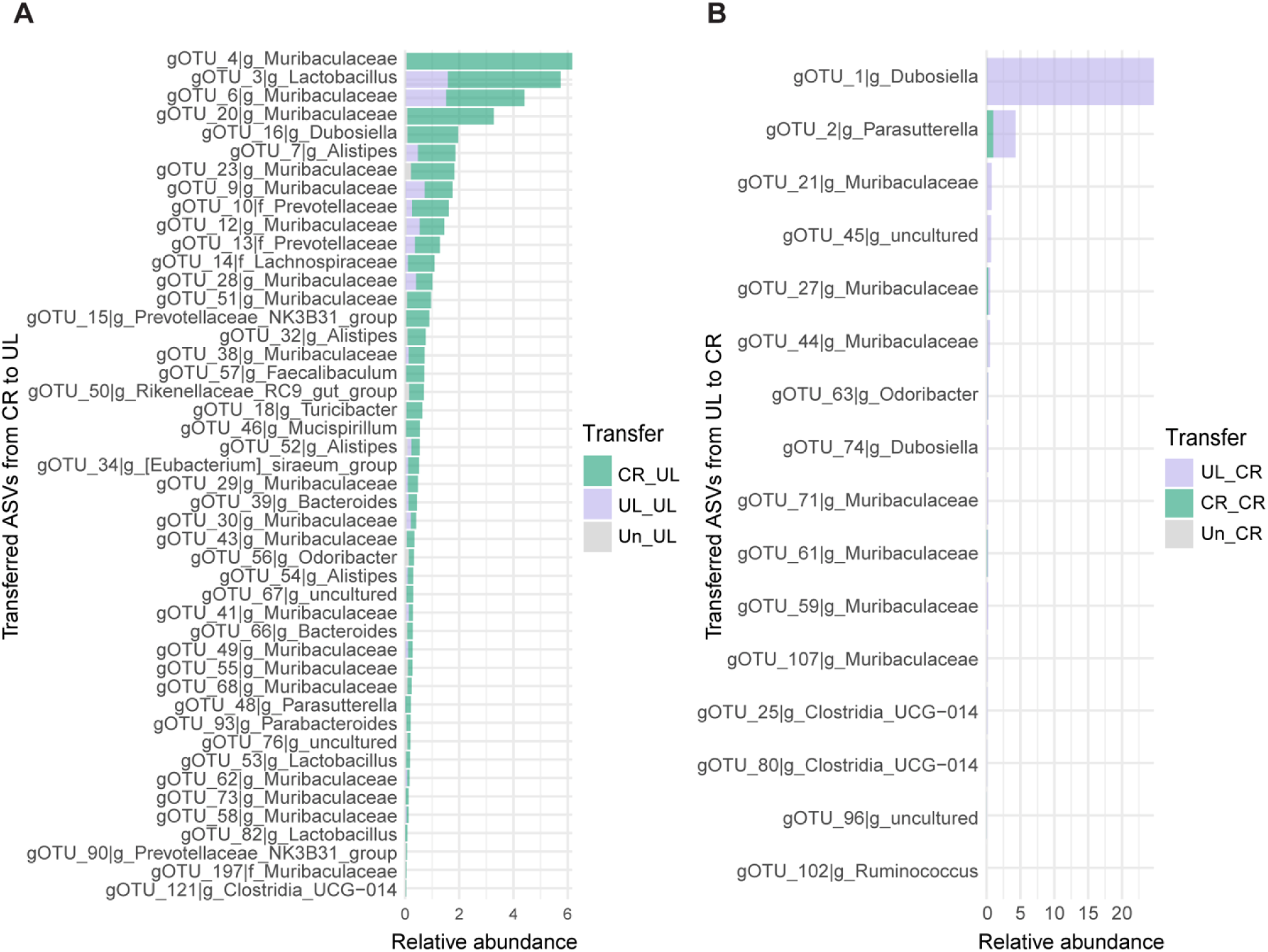
Significantly transferred gut OTUs during co-housing. (A) Gut OTUs significantly transferred from CR mice in the co-housed groups (Mix_CR; n = 14) to the UL mice (Mix_UL; n =16) during 28 days of co-housing; (B) Gut OTUs significantly transferred from Mix_UL mice to Mix_CR mice during co-housing. Different colours represent different source_sink combinations: CR_UL refers to transferring from CR mice to UL mice; UL_UL represents inheritance from UL mice between day -28 and day 0; Un_UL indicates the contribution from unknown environments to UL mice after co-housing; UL_CR refers to transferring from UL mice to CR mice; CR_CR represents inheritance from CR mice between day -28 and day 0; Un_CR indicates the contribution from unknown environments to CR mice after co-housing. The bars indicate the mean relative abundance of gut OTUs being transferred. The proportion of each gut OTU in each sink sample after co-housing from different source environments before co-housing was calculated with Sourcetracker. The transferred relative abundance was calculated based on the original relative abundance in each sink sample (day 0) and mean abundance of transferred OTU in each group was calculated and compared Wilcoxon test, and p-values were adjusted with the Benjamini-Hochberg method.

Next, we assessed the compositional dynamics of the skin microbiota during co-housing, performing the same analyses as conducted for the gut microbiota. However, unlike the gut microbiota, the skin microbiota did not exhibit a similarly pronounced homogenization. Specifically, significant differences in the Shannon index were observed between Mix_CR and Mix_UL mice both before and after co-housing. Furthermore, beta diversity analysis reveals marginally significant differences before-, and significant differences after co-housing (**Fig. S4A-C**). Despite the lack of homogenization based on assessment by beta diversity, 29 OTUs were identified as exchanged during co-housing using SourceTracker, with 27 transferred from Mix_CR to Mix_UL and 2 transferred from Mix_UL to Mix_CR (**Fig. S4D-E; Supplementary Table 4**). Among the OTUs transferred from Mix_CR to Mix_UL, sOTU_219 (matching the genus *Deinococcus*) was reported to be negatively correlated with skin inflammatory diseases, such as psoriasis and allergic skin inflammation (49). Additionally, sOTU_1058 (*Staphylococcus epidermidis*), which was enriched in CR mice prior to PD induction, was also identified among the transferred OTUs (**Fig. S4D-E & Fig. S2B**). Thus, altogether the skin microbiota also changed during co-housing, and although to a lesser degree, the asymmetric pattern of much more Mix_CR to Mix_UL transfer compared to Mix_UL to Mix_CR transfer is consistent with the observations for the gut microbiota.

### 3. The effect of PD induction on gut and skin microbiota

Fecal samples and skin punch biopsies collected on day 12 allowed us to assess the effects of PD induction using two approaches: (i) cross-sectional comparison of BP-like EBA and NR groups at day 12, and (ii) longitudinal comparison within the BP-like EBA group between baseline (day 0) and post-induction (day 12). Beta diversity analysis revealed significant differences between PD and NR groups for both gut (p = 0.003) and skin microbiota (p = 0.001), while alpha diversity changes were limited, with the exception of a reduced Chao1 index in the gut microbiota of UL mice (**Fig. 6A-D**). LEfSe analysis (p.adj < 0.05, LDA > 2) identified group-specific shifts in taxa. In UL mice, three gut OTUs are significantly reduced in the PD group, including two *Faecalibaculum* OTUs, one corresponding to *F. rodentium*, a regulator of epithelial homeostasis (50), and gOTU_22 (*Ligilactobacillus murinus*; NCBI, E-value = 3e-158, identity = 100%), known to mediate anti-inflammaging effects in calorie-restricted mice (51). In CR mice, four gut OTUs decrease, including gOUT_50 (*Alistipes putredinis*; NCBI, E-value = 3e-138, identity = 97.32%), a taxon associated with reduced inflammation (52). No differentially abundant gut OTUs are detected in co-housed groups (Mix_UL and Mix_CR) (**Fig. 6E; Supplementary Table 5**). For the skin microbiota, significant differences are observed only in CR mice, where 10 OTUs changed in abundance. Notably, sOTU_1058 (*S. epidermidis*) increased in the PD group (**Fig. 6F; Supplementary Table 5**).

**Fig. 6.**
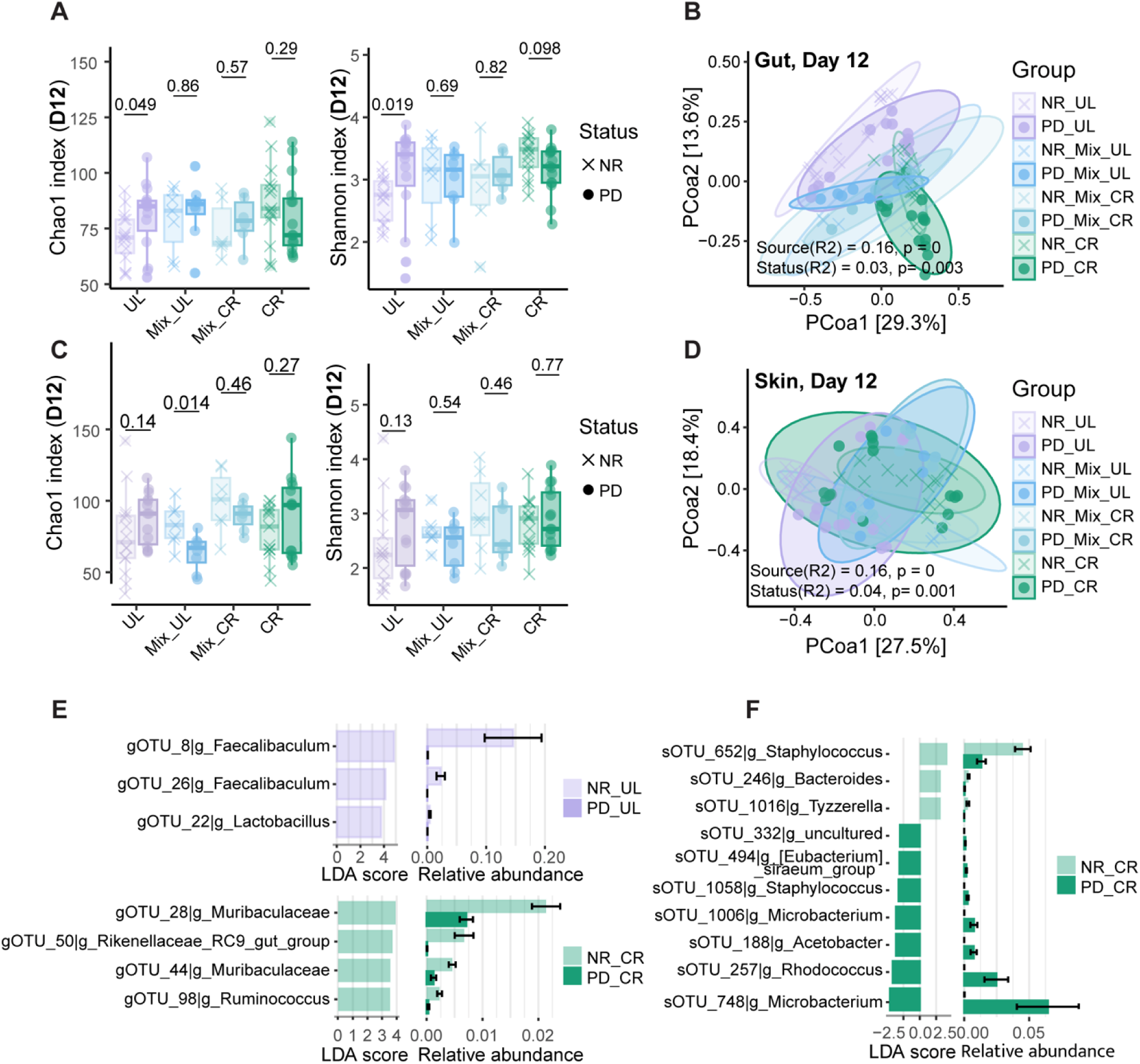
Effects of PD induction on the gut and skin microbiota. (A-B) Alpha diversity and beta diversity of gut microbiota in mice injected with anti-COL7 IgG (PD groups) and those injected with normal rabbit IgG as controls (NR groups) on day 12, the endpoints of PD induction; (C-D) Alpha diversity and beta diversity of skin microbiota in mice injected with anti-COL7 IgG (PD groups) and those injected with normal rabbit IgG as controls (NR groups); (E) Differential gut OTUs were identified in mice between PD and NR groups on day 12; (F) Differential skin OTUs were identified in mice between PD and NR groups on day 12. The alpha indices were compared with Wilcoxon test. Beta diversity was analysed using PERMANOVA with 9,999 permutations. The statistical p values for differential OTUs were adjusted by the Benjamini-Hochberg procedure, and the OTUs with LDA > 2 and adjusted p value < 0.05 were considered as significantly differential.

Longitudinal comparison of baseline (day 0) and post-induction (day 12) samples within PD groups (CR, UL, Mix_CR, Mix_UL) revealed that the gut microbiota of CR mice remained relatively stable. Significant beta diversity changes were detected only in UL mice, whereas Mix_UL, Mix_CR, and CR mice showed no significant shifts. The increasing p-values for these latter three groups (Mix_UL < Mix_CR < CR) suggest a gradient of stability, with CR mice being the most stable (**Fig. S5A**). Differential abundance analysis identified only one significantly decreased gut taxon, gOTU_22, in the PD-UL group (p.adj < 0.05, LDA > 2) (**Fig. S5B**). For the skin microbiota, significant beta diversity changes were observed in all eight groups, including both NR and PD mice from each source, indicating that environmental factors, in addition to PD induction, influenced community composition over the 12-day period (**Fig. S6A**). Differentially abundant skin OTUs (p.adj < 0.05, LDA > 2) that changed exclusively in PD groups were considered potentially BP-like EBA-associated. For example, sOTU_166 (*Acinetobacter*) decreased in PD_UL and PD_CR groups, whereas sOTU_181 (*Lachnospiraceae_NK4A136_group*) increased in PD_UL and PD_Mix_UL mice (**Fig. S6B–C; Supplementary Table 6**).

### 4. Prediction of PD severity from baseline gut and skin microbiota

Given the observed associations between baseline microbiota composition and subsequent disease outcomes, we first tested for correlations between the relative abundance and diversity metrics of gut and skin microbiota prior to PD induction and PD severity scores on day 12. Only one feature, gut gOTU_67, shows a significant association, displaying a negative correlation with disease severity (corr = −0.61, p.adj = 0.0197) after adjusting for mouse source (**Fig. 7A**).

**Fig. 7.**
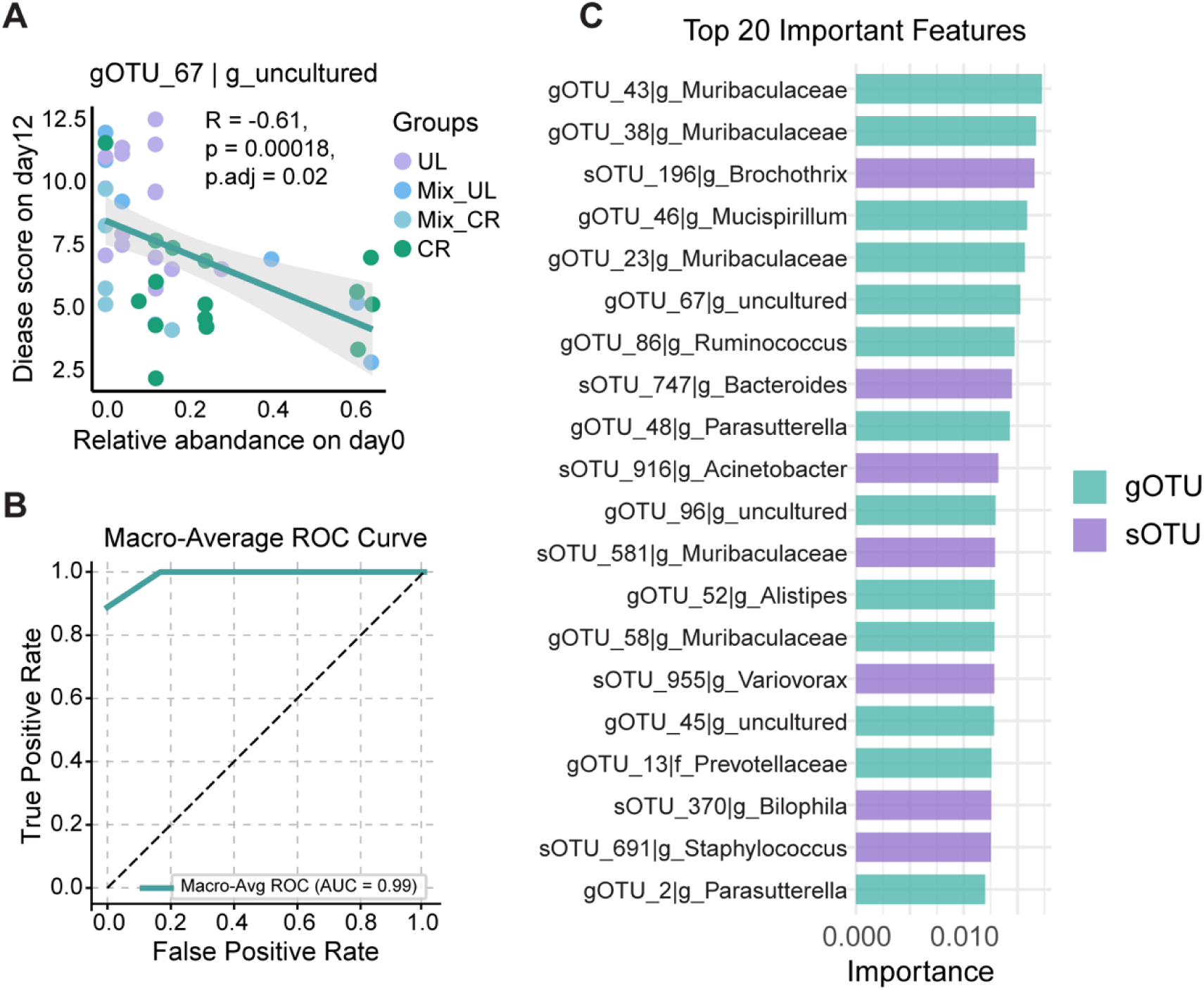
Prediction potential of the gut and skin microbiota before induction for PD severity. (A) The only gut OTU significantly correlated with PD severity. The correlation between abundance of gut and skin OTUs before induction were calculated with PD severity of all 45 mice after adjusting sources (CR, UL and Mix_CR and Mix_UL). The statistical p values were adjusted by the Benjamini-Hochberg procedure; (B) The Macro-average Receiver Operating Characteristic (ROC) curve of the multi-class XGBClassifier (XGB). The curve shows the averaged true positive rate (TPR) across all classes at varying false positive rates (FPRs), with each class evaluated as a one-vs-rest task. The high macro-average AUC (0.99) indicates strong model performance across all classes, independent of class imbalance. Dashed line represents random chance (AUC = 0.5); (C) The 20 most important OTUs contributed to the optimized XGBClassifier model which can well classify the disease severity levels.

To further evaluate the predictive potential of the microbiota, we applied a machine learning approach integrating both gut and skin microbial features. Disease severity was first categorized into three levels (low (2.715-5.25), medium (6.625-8.2625), and high (9.225-12.5)), and XGBClassifier, a widely used gradient boosting algorithm, was trained on the combined dataset. Data were randomly split into training (80%) and testing (20%) sets, and key hyperparameters were optimized using 5-fold cross-validation, yielding a final model with a learning rate of 0.01, maximum tree depth of 6, and 50 trees. The optimized model achieved strong predictive performance, with an accuracy of 88.89%, a macro F1-score of 88.57%, and a macro-average AUC of 99% for the multiclass task (**Fig. 7B**).

Feature importance analysis revealed that gut microbiota contributed more to classification performance than skin microbiota (53.64% vs. 46.36%), with 69 of 112 gut OTUs and 63 of 161 skin OTUs identified as important predictors. Gut OTUs dominated the top 20 most important features (13/20), including gOTU_67, gOTU_43, and gOTU_38 (**Fig. 7C; Supplementary Table 7**). Notably, gOTU_67, which we also detected as transferred from Mix_CR to Mix_UL during co-housing, emerged as the strongest single predictor.

To investigate its potential functional role, we aligned the representative sequence of gOTU_67 against multiple 16S rRNA gene databases (NCBI, Silva v138 (53) and GTDB (54)). The closest match was *Scatocola sp*. GCA_910577205.1 in GTDB (99.4% nucleotide identity over 205 aligned positions) (**Fig. S7A**). The representative genome (completeness 100%, contamination 0.3%; CheckM2 (55)) was functionally annotated using Prokka, revealing enrichment in pathways related to translation, ribosomal structure and biogenesis, cell wall/membrane biogenesis, and carbohydrate transport and metabolism (COG analysis; **Fig. S7B**) (56, 57). CAZyme annotation identified GT2 and GT4 (glycosyltransferases), GH23 (glycoside hydrolase), and CBM50 (carbohydrate-binding module) as the most abundant carbohydrate-active enzymes (**Fig. S7C**) (58). This suggests a capacity for complex carbohydrate metabolism, which could e.g. influence gut barrier function and/or immune modulation.

## Discussion

Overall, our study provides important new evidence that the resident microbiota can influence the onset and progression of experimental PD in mice. We observe that initial differences in microbiota composition, particularly in the gut, among mice from different sources (UL, CR, and co-housed groups) are associated with variation in BP-like EBA severity. Co-housing experiments demonstrate active microbial exchange, with asymmetric transfer from CR to UL mice. This results in decreased disease severity in co-housed UL mice, suggesting that the introduction of protective taxa can modulate disease outcomes. Finally, the predictive potential of microbial features, particularly those derived from the gut, highlights the possibility of using them as biomarkers for disease outcome.

Given that the groups of mice we analysed share the same genetic background, the differences in disease severity argue that host-specific environmental factors play an important role in disease susceptibility, with microbiota composition as the likely key contributor. UL mice developed significantly more severe disease than CR mice, while co-housed mice showed intermediate severity. Notably, co-housed CR mice exhibited disease levels comparable to CR mice, consistent with a protective effect from microbes transferred from Mix_CR to Mix_UL.

Baseline gut and skin microbial diversity also differed significantly between CR and UL mice (Shannon index and beta diversity). For the gut, the Shannon index showed a trend of an inverse correlation with disease severity across groups, with CR mice having the highest diversity. Although not statistically significant (Chao1: r = −0.12, p = 0.44; Shannon: r = −0.09, p = 0.58), the pattern suggests that greater gut microbial diversity may confer protection against inflammation, potentially contributing to the lower disease severity in CR mice. Similarly, skin microbiota diversity (Shannon index) was significantly higher in CR mice, which is consistent with a previous study of experimental BP-like EBA that linked greater pre-induction diversity to the absence of disease symptoms after induction (59). Differential abundance analysis further identifies the enrichment of anti-inflammatory and immunomodulatory taxa in CR mice, including *L. intestinalis* and *P. distasonis* (38, 39). In contrast, UL mice were enriched in taxa linked to pro-inflammatory responses, such as *T. muris* and *M. intestinale* (34, 35). Together, these findings underscore the influence of microbiota composition on the extend of skin disease outcomes and highlight the potential protective role of taxa present in the CR gut microbiota.

Co-housing CR and UL mice led to partial homogenization of the gut microbiota, with Mix_UL and Mix_CR communities becoming more similar to each other than to their original sources. This convergence was accompanied by an increase in gut microbial diversity in UL mice, suggesting a net gain of taxa from CR donors. Sourcetracker analysis confirms an asymmetric exchange, with substantially more OTUs transferred from CR to UL mice (46 vs. 16 in the opposite direction). Many of these transferred taxa, including *L. johnsonii*, *Alistipes spp*., and *Prevotellaceae_NK3B31_group*, are known for anti-inflammatory or barrier-protective functions (36, 37, 44–47), and their introduction may explain the intermediate disease severity observed in Mix_UL mice. In contrast, skin microbiota composition was less affected by co-housing. Although some taxa such as *S. epidermidis* were exchanged, the overall pattern of beta diversity remained more source-dependent, suggesting greater resilience of skin microbial communities compared to the gut. This differential responsiveness provides evidence that the gut microbiota plays a more central role in modulating PD severity in our model.

PD induction induced significant changes in both gut and skin microbiota, with distinct responses observed among UL, CR, and co-housed mice, based on comparisons between PD and NR groups on day 12 as well as the longitudinal comparison between day 0 and day 12 within PD groups. The gut microbiota of CR mice remains relatively stable, while UL mice exhibit changes in beta diversity and a reduction in potentially beneficial taxa such as *F. rodentium,* and *L. murinus*. These changes may contribute to the higher PD disease severity observed in UL mice. In contrast, the skin microbiota showed significant changes across all groups, suggesting that environmental factors, in addition to PD induction, influence skin microbial composition. Notably, *S. epidermidis* (sOTU_1058) increased in the BP-like EBA group for CR mice. Given its reported ability to suppress pro-inflammatory responses and enhance colonization resistance against *S. aureus* (43), this expansion may contribute to the relative protection observed in CR mice.

In addition to diversity patterns, taxa-level shifts, and microbial exchange, our predictive modeling highlights gut gOTU_67 as a strong indicator of PD disease severity. gOTU_67 negatively correlates with disease severity, dominates the feature importance rankings in the XGBClassifier model, and is among the taxa transferred from CR to UL mice during co-housing. Genomic annotation of its closest match, *Scatocola sp.*, reveal capacities for carbohydrate metabolism, cell wall biogenesis, and ribosomal function, suggesting potential roles in maintaining gut barrier integrity, modulating immune responses, or competing with pro-inflammatory taxa. These traits provide plausible mechanistic hypotheses for its association with reduced disease severity, making it a promising candidate micro-biomarker. Future isolation and characterization of *Scatocola sp.* may help determine its preventative and/or therapeutic potential for modulating autoimmune skin disease outcomes.

In conclusion, this study experimentally demonstrates that the host-associated microbiota can influence the severity of experimental PD. Baseline microbiota composition in the gut in particular is linked to disease outcomes, and co-housing experiments revealed that the transfer of microbes from CR to UL mice reduced disease severity. While the gut microbiota partially homogenized during co-housing, the skin microbiota remained largely source-dependent, suggesting greater resilience to environmental change. Machine learning identified *Scatocola sp.* as a strong predictor of disease severity. These findings lay the groundwork for future studies using gnotobiotic or microbiota-depleted models. Ultimately, microbiota-based interventions— such as targeted probiotics, fecal microbiota transplantation, or species-specific phage therapy, may serve as promising future strategies to prevent or mitigate PD.

## Materials and methods

### Mice

A total of 90 female C57BL/6J mice (8-14 weeks old at the starting timepoint of co-housing experiment) were sourced from two breeding sources: University of Lübeck (UL, n = 46) and Charles River Laboratories (CR; Sulzfeld, Germany, n = 44). Among those mice, 16 from UL and 14 CR were randomly chosen and co-housed for 28 days, with 5 in each cage, with a dark-light cycle of 12h:12h and under a constant temperature and humidity. All mice were maintained under specific pathogen-free (SPF) conditions with *ad libitum* access to standard water and chow. All protocols were approved by the former ministry for energy transition, agriculture environment, nature and digitalization Schleswig-Holstein (41-5/18).

### Passive BP-like EBA mouse model induction

The passive BP-like EBA mouse model was induced in mice by injection of anti-COL7 IgG, with age-matched controls receiving normal rabbit IgG. Injections were administered every two days over a 12-day course. Disease severity was assessed based on the extend of clinical manifestations, including crusts, erythema, lesions, and/or alopecia on individual body parts on day 4, 8, and 12 according to an established scoring system (60).

### Sample collection

Fecal samples and skin punch biopsies were collected from mice at three timepoints: (1) starting of co-housing, (2) prior to disease induction, and (3) experimental endpoint. For fecal sampling, fresh pellets were aseptically collected and immediately frozen at -80°C until further use. Skin microbiota sampling was performed on punch biopsies of ear tissue as previously described (31).

### Nucleic acid extraction

Fecal DNA was extracted using QIAamp PowerFecal Pro DNA Kit (Qiagen, Cat. No. 51804) according to the manufacturer’s protocol. For skin-derived RNA extraction, punch biopsies were extracted using the AllPrep DNA/ RNA 96 Kit (Qiagen, Cat. No. 80311) as previously described (61, 62). All extracted DNA and RNA samples were stored at -80°C until further use.

### 16S rRNA gene sequencing and data processing

The V1-V2 region of the 16S rRNA gene was amplified using 27F and 338R primers using a dual barcoding approach as previously described (63). The obtained library was subsequently sequenced on an Illumina Miseq sequencer (250PE). Demultiplexing was performed using the Casava pipeline (Illumina), retaining only barcodes with zero mismatches. Raw 16S rRNA gene sequencing data were processed using QIIME2 (v2022.8) (64). Sequences were denoised using DADA2 (65), trimming the first 7 (forward) or 9 (reverse) bases from the 5’ end and truncating at 230 (forward) or 200 (reverse) bases from the 3’ end. Reads were also truncated at the first instance of a quality score ≤ 3. The denoised sequences were then clustered at 99% similarity OTUs. An abundance table was generated, and taxonomic annotation was performed using the SILVA 138 database (53).

To control for the influence of contamination in low-biomass skin microbiota, we included negative controls for sampling environment, RNA extraction and library preparation. We then identified potential contaminants with the “prevalence” method with a threshold of in the R package “decontam” (R version: 4.2.2; decontam version: 1.18.0), and 236 potential contaminants were removed out of a total of 7767 OTUs. Moreover, the sequences which belong to families *Halomonadaceae* or *Shewanellaceae* (n = 5) were removed as they are deemed common contaminants in low-biomass samples (66). Further, we applied an advanced normalization method to control for noise in skin microbiota data, as described previously (61). In brief, we randomly subsampled 2,900 reads (the minimum depth among skin samples) 1,000 times for each sample. We then examined the distribution of OTU frequencies across these 1,000 subsamples. OTUs whose frequency met or exceeded the 10th percentile of the overall frequency distribution were retained, and a final subsample of 2,900 reads was drawn exclusively from this set of selected OTUs. For gut microbiota, the samples were randomly rarefied to the minimum read count of 2,500.

For OTUs of interest, the representative sequences (ASVs) were further aligned in the NCBI database. If no reliable match was found in NCBI (e.g., for gut gOTU_67), we extended the alignment to the 16S rRNA bac120 representative sequences in the Genome Taxonomy Database (GTDB) using the USEARCH global alignment tool (54, 67). The representative genome matched to the gut gOTU_67 was obtained from GTDB, and genes were then predicted from the representative genome with prokka and annotated using Clusters of Orthologous Groups of proteins (COG) and Carbohydrate-Active enZYmes (CAZymes) databases (56–58).

### Microbial diversity

Diversity analyses were performed using the R package vegan (R v4.2.2; vegan v2.6-4). Alpha diversity (Chao1 and Shannon indices) was compared between groups using the Wilcoxon test. Beta diversity, based on Bray-Curtis dissimilarity, was analysed using PERMANOVA (68) with 9,999 permutations via the “adonis” function.

### Disease-associated gut and skin microbial OTUs identification

Linear discriminant analysis effect size (LEfSe) analysis (69) was used to identified differential gut and skin microbial OTUs with an absolute value of LDA score no less than 2, and an adjusted p value <0.05. The LEfSe was performed using R package microeco (v 1.4.0). Spearman correlations between the abundance of gut and skin OTUs and disease severity were calculated in all 45 mice after adjusting sources (CR, UL and Mix_CR and Mix_UL) using the “cor.test” function in R. To avoid the influence of zero-inflation, only species or pathways with a prevalence of at least 20% were selected to be included in the correlation analysis. When appropriate, statistical p values were adjusted by the Benjamini-Hochberg procedure.

### Machine learning analysis of microbiota and PD disease severity

To explore the potential of microbiota in characterizing PD disease severity, we constructed machine learning models using microbial features from gut and skin data. Disease severity was categorized into three levels, and the XGBClassifier (XGB) alglorithm was applied to the integrated dataset. The data were randomly split into training and testing sets (80:20 ratio) for 100 independent iterations to ensure robust evaluation. Hyperparameters were optimized using 5-fold cross-validation, resulting in a final model with a learning rate of 0.01, a maximum tree depth of 6, and 50 trees. The optimized model achieved an accuracy of 88.89% and a PRAUC of 97.22%. Feature importance analysis was conducted to identify key microbial features contributing to disease severity classification. The analysis was performed with xgboost model (v 2.1.4) in python (v 3.9.13).

### Microbial Community Transfer Analysis

To access microbial transfer between co-housed individuals, we applied Sourcetracker, a Bayesian-based algorithm (48). The microbial communities after co-housing (Mix_UL_D0 and Mix_CR_D0) were designated as sink environments, while profiles before co-housing (Mix_UL_Dm28 and Mix_CR_Dm28) served as source environments. For each sink sample, the proportion of OTUs derived from different sources was estimated. These proportions were integrated, and transferred relative abundances were calculated based on the original abundance of each OTU. Finally, the mean abundance of transferred OTUs in each group was compared using the Wilcoxon test, with p-values adjusted by the Benjamini–Hochberg method.

## Data Availability

The data presented in this study are openly available in the NCBI SRA database under BioProject accession number PRJNA1315180 at https://www.ncbi.nlm.nih.gov/bioproject/. Reviewer link: https://dataview.ncbi.nlm.nih.gov/object/PRJNA1315180?reviewer=m7sskeephnfnoadq9b53891 oe0.

## Acknowledgements

We thank Katja Cloppenborg-Schmidt for excellent technical assistance and Sven Künzel for sequence data generation. ChatGPT5 (OpenAI) was used for the improvement of originally composed text.

## Author Contributions

Conceptualization, S.I., E.S. and J.F.B. Data analysis, X.L., S.P., Y.M., A.C. Investigation, S.P., K.B., H-O.D. Writing – Original Draft Preparation, X.L., Y.M., J.F.B. Writing – Review & Editing, all authors; Visualization, X.L., Y.M., A.C., J.F.B.; Supervision, M.P., M.G., S.I., E.S. and J.F.B. Project Administration S.I., E.S. and J.F.B. Funding Acquisition, S.I., E.S. and J.F.B.

