## Supplementary Figures for "Gut Microbiota Modulates and Predicts Disease Severity in Experimental Pemphigoid Disease"

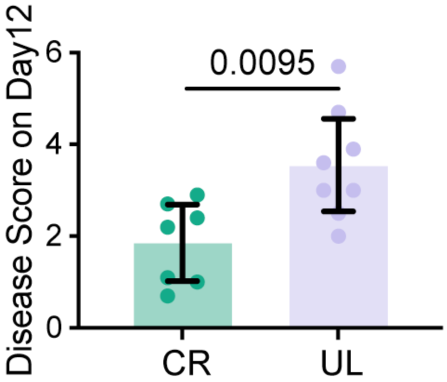


**Fig. S1 Evaluation of disease severity of PD mice induced from mice with different breeding sources in the preliminary experiment.** The disease score for mice from CR (n = 7) and UL (n = 8) sources mice on day 12. The scores were compared with t test. The different colours represent different groups.

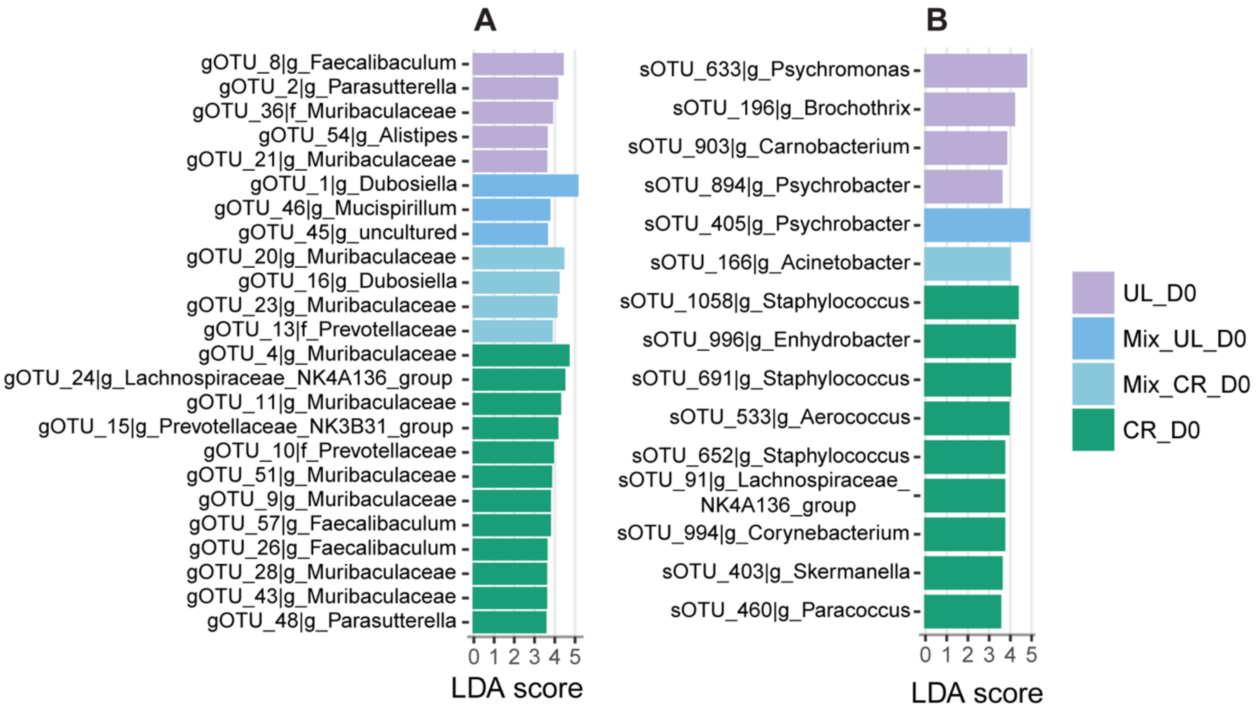


**Fig. S2 Differential gut and skin OTUs of mice from different sources identified before PD induction.** (A) Differential gut OTUs were identified in mice from different sources on day 0, the starting day of PD induction, with Linear Discriminant Analysis Effect Size (LEfSe) analysis; (B) Differential skin OTUs were identified in mice from different sources on day 0 with LEfSe analysis. The statistical p values were adjusted by the Benjamini-Hochberg procedure, and the OTUs with LDA > 2 and adjusted p value < 0.05 were considered as significantly differential, with only presenting differential OTUs with LDA > 3.5.

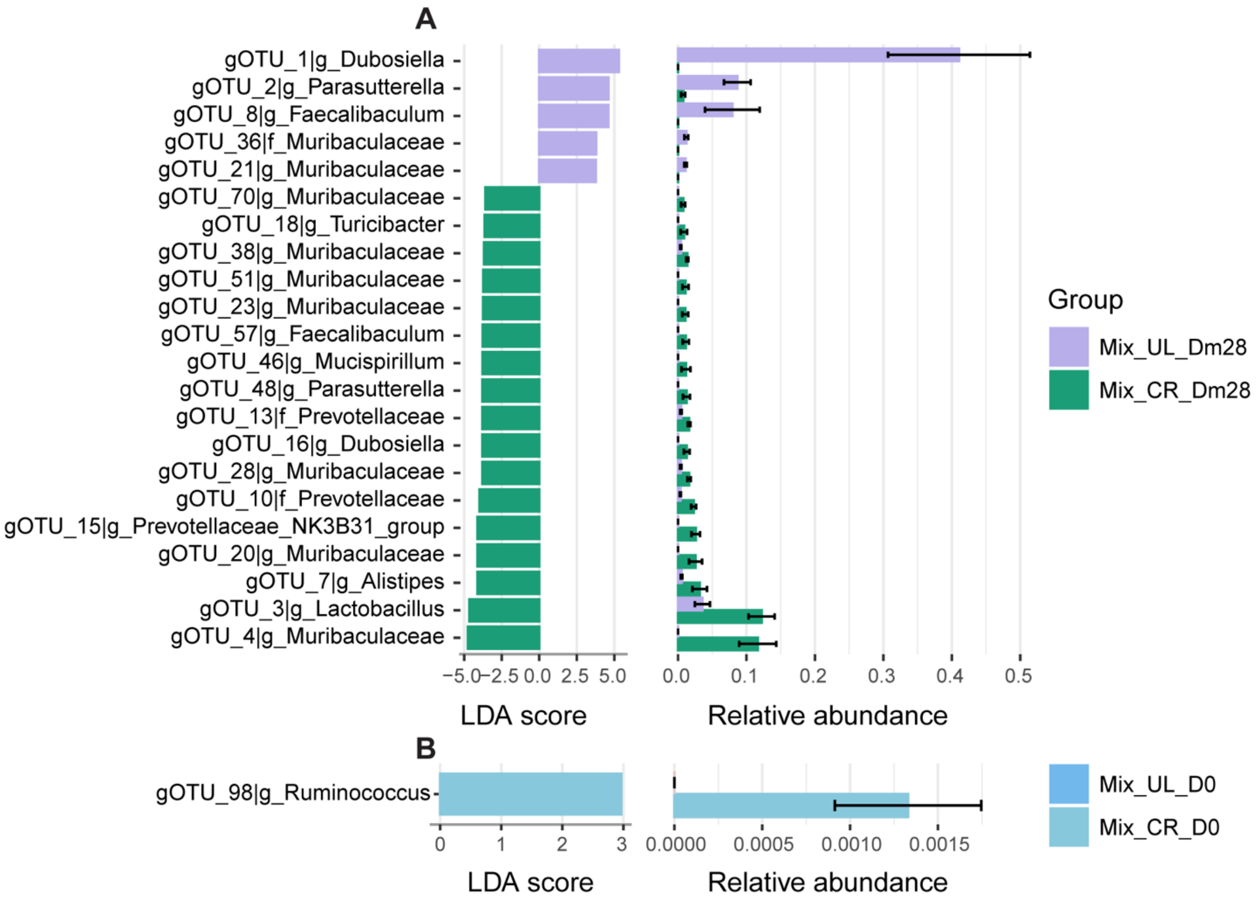


**Fig. S3 Differential gut OTUs of mice between co-housed UL and CR mice before and after co-housing.** (A) Differential gut OTUs before co-housing (Dm28) were identified in co-housed mice with LEfSe analysis; (B) Differential gut OTUs after co-housing (D0) were identified in co-housed mice. The statistical p values were adjusted by the Benjamini-Hochberg procedure, and the OTUs with LDA > 2 and adjusted p value < 0.05 were considered as significantly differential.

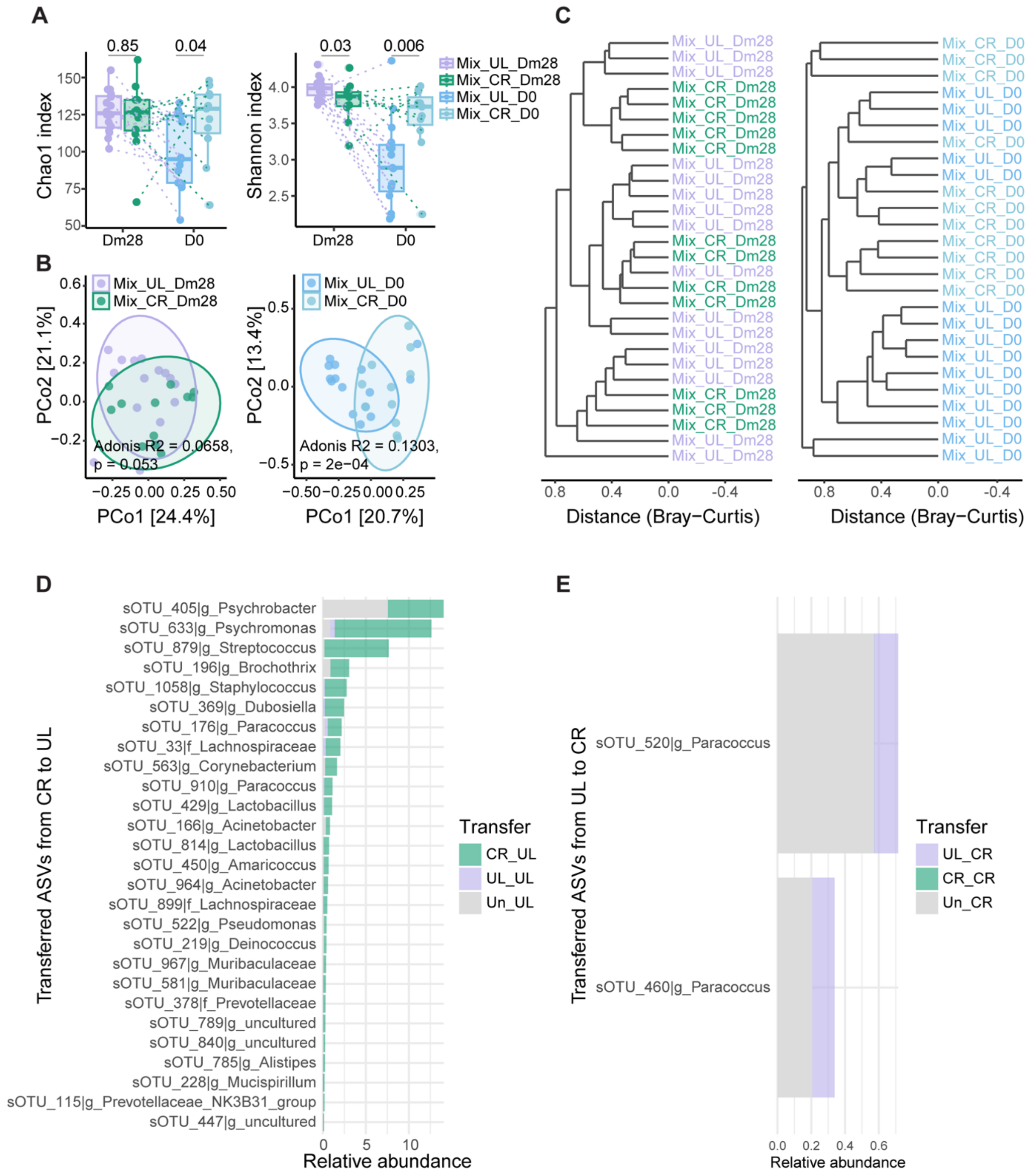


**Fig. S4 Co-housing effects on skin microbiota.** A) Chao1 and Shannon indices of skin microbiota in 30 co-housed mice before (Dm28) and after (D0) co-housing, with lines pairing the samples from same individual mouse; (B) The PCoA plots of skin microbiota of co-housed UL and CR mice before (Dm28) and after (D0) co-housing; (C) The hcluster plots of skin microbiota on day -28 and day 0. (C) Significantly transferred skin OTUs during co-housing. The alpha indices and mean abundance of transferred OTU were compared with Wilcoxon test. Beta diversity was analysed using PERMANOVA with 9,999 permutations.

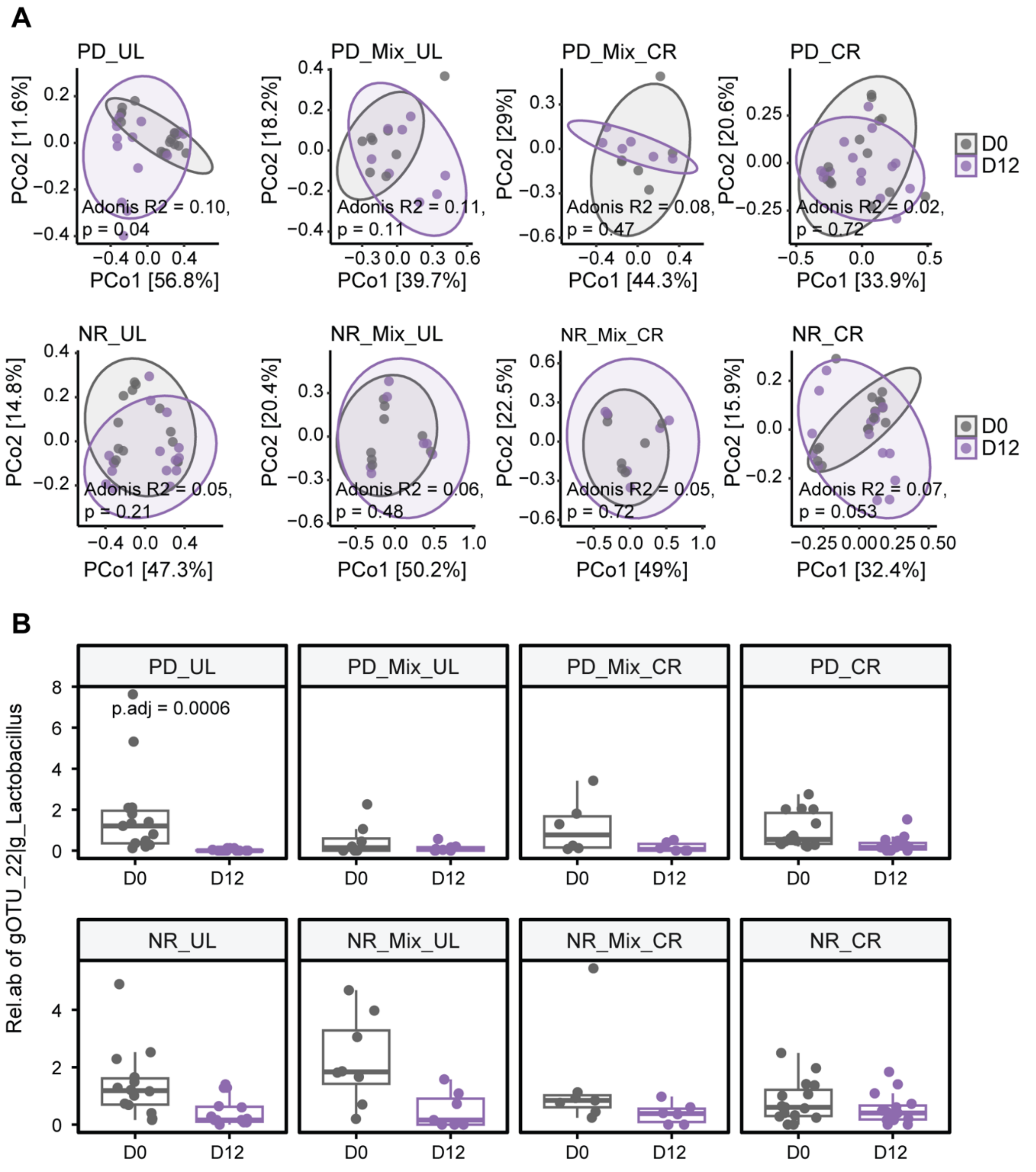


**Fig. S5 Longitudinal effects of PD induction on gut microbiota.** (A) Beta of gut microbiota in BP-like EBA mice before (D0) and after PD induction (D12); (B) The relative abundant of the only differential gut OTUs identified with LEfSe analysis after PD induction in UL mice. Beta diversity was analysed using PERMANOVA with 9,999 permutations. The statistical p values for differential OTUs were adjusted by the Benjamini-Hochberg procedure, and the OTUs with LDA > 2 and adjusted p value < 0.05 were considered as significantly differential.

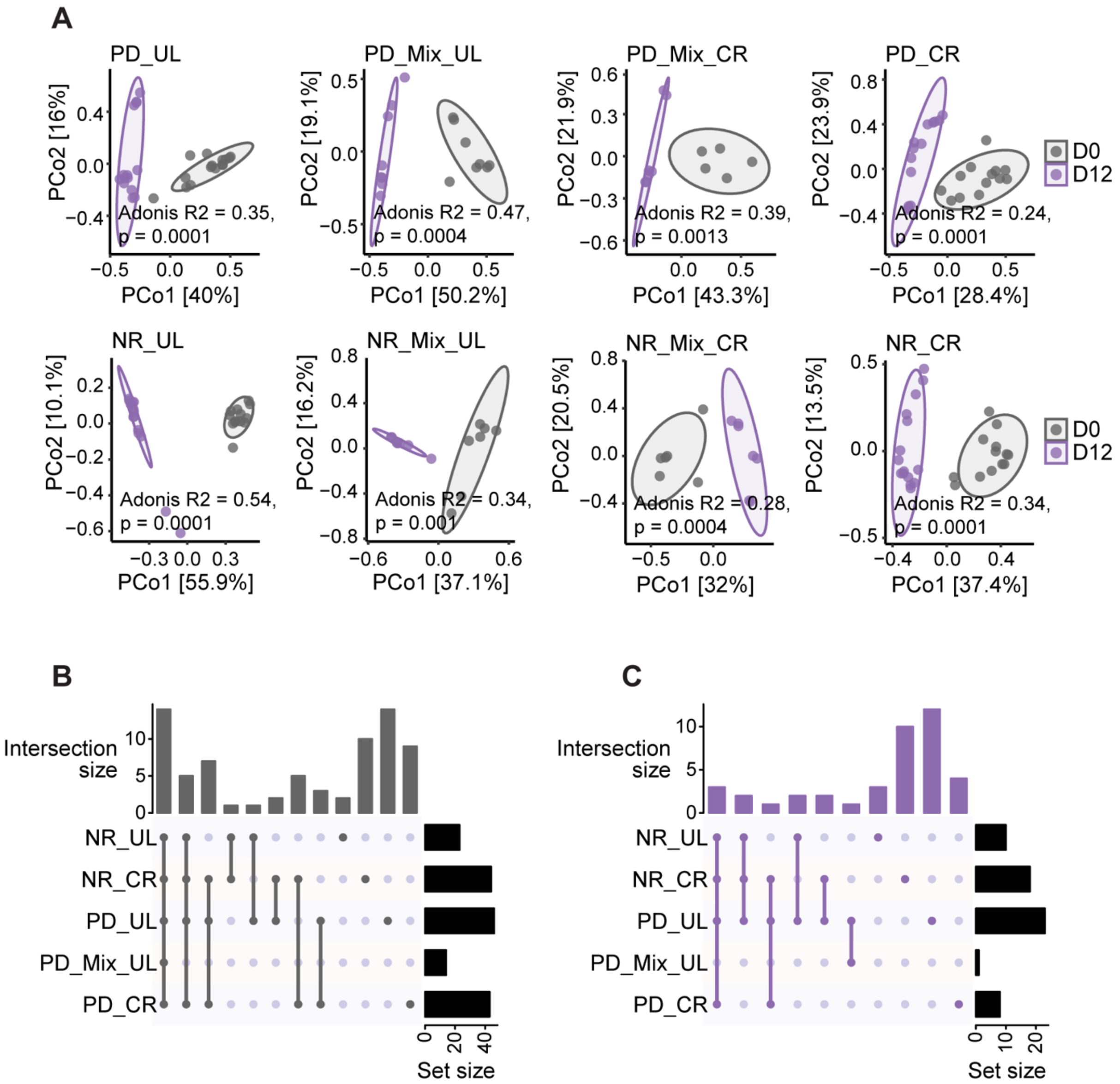


**Fig. S6 Longitudinal effects of PD induction on skin microbiota.** (A) Beta of skin microbiota in PD mice before (D0) and after PD induction (D12); (B) The UpSet plots showing the distribution significantly changed skin OTUs after 12 days of PD induction in mice, which were identified with LEfSe analysis. Beta diversity was analysed using PERMANOVA with 9,999 permutations. The statistical p values for differential OTUs were adjusted by the Benjamini-Hochberg procedure, and the OTUs with LDA > 2 and adjusted p value < 0.05 were considered as significantly differential.

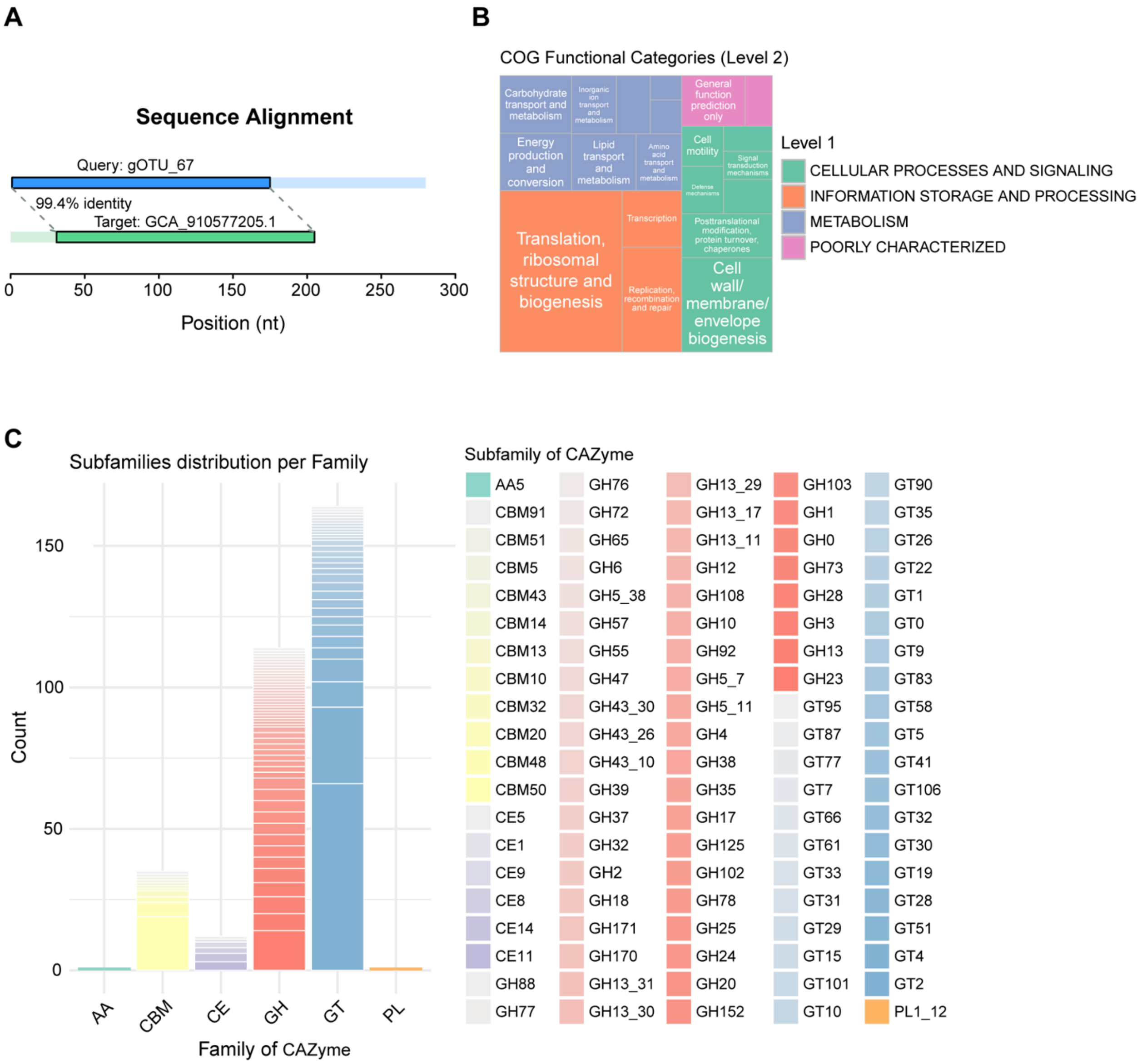


**Fig. S7 Taxonomic classification and functional potential of gut-derived OTU_67 based on GTDB genomic alignment.** (A) Sequence alignment of gut OTU_67 (query) with the Genome Taxonomy Database (GTDB) 16S rRNA bac120 representative sequence of *Scatocola sp.* GCA_910577205.1 (reference) revealed 99.4% nucleotide identity over 205 aligned positions (USEARCH global alignment, -id 0.97). (B) Functional categorization of predicted genes based on the Clusters of Orthologous Groups (COG) database, showing enrichment in information processing (especially translation, ribosomal structure and biogenesis), cellular processes (e.g., cell motility, cell wall biogenesis). (C) Predicted carbohydrate-active enzymes (CAZymes) assigned to gOTU_67, grouped by major families: auxiliary activities (AA), carbohydrate-binding modules (CBM), carbohydrate esterases (CE), glycoside hydrolases (GH), glycosyltransferases (GT), and polysaccharide lyases (PL). The barplot represents the number of predicted genes per subfamily.
